# Decision support for managing an invasive pathogen through efficient clean seed systems: Cassava mosaic disease in Southeast Asia

**DOI:** 10.1101/2024.02.13.580210

**Authors:** Kelsey F. Andersen Onofre, Erik Delaquis, Jonathan C. Newby, Stef de Haan, Thuy Cu Thi Le, Nami Minato, James P. Legg, Wilmer J. Cuellar, Ricardo I. Alcalá Briseño, Karen A. Garrett

## Abstract

**CONTEXT:** Effective seed systems must both distribute high-performing varieties efficiently and slow or stop the spread of pathogens and pests. Epidemics increasingly threaten crops around the world, endangering the incomes and livelihoods of smallholder farmers. Responding to these food and economic security challenges requires stakeholders to act quickly and decisively during the early stages of invasions, typically with very limited resources. The recent introduction of cassava mosaic virus into southeast Asia threatens cassava production in the region.

**OBJECTIVES:** Our goal in this study is to provide a decision-support framework for efficient management of healthy seed systems, applied to cassava mosaic disease. The specific objectives are to (1) evaluate disease risk in disease-free parts of Cambodia, Lao PDR, Thailand, and Vietnam by integrating disease occurrence, climate, topology, and land use, using machine learning; (2) incorporate this predicted environmental risk with seed exchange survey data and whitefly spread in the landscape to model epidemic spread in a network meta-population model; and (3) use scenario analysis to identify candidate regions to be prioritized in seed system management.

**RESULTS AND CONCLUSIONS:** The analyses allow stakeholders to evaluate strategy options for allocating their resources in the field, guiding the implementation of seed system programs and responses. Fixed rather than adaptive deployment of clean seed produced more favorable outcomes in this model, as did prioritization of a higher number of districts through the deployment of smaller volumes of clean seed.

**SIGNIFICANCE:** The decision-support framework presented here can be applied widely to seed systems challenged by the dual goals of distributing seed efficiently and reducing disease risk. Data-driven approaches support evidence-based identification of optimized surveillance and mitigation areas in an iterative fashion, providing guidance early in an epidemic, and revising them as data accrue over time.

## Introduction

Seed systems must play the dual role of effectively distributing good crop varieties and limiting the dispersal of crop pathogens and pests. Invasive plant pathogens are posed to inflict ever-greater damage as planted area and density of field crops increase globally (Ristaino et al., 2021). Fueled by increasingly globalized trade, growing cropland connectivity, and the impacts of climate change, crop pest and disease outbreaks have been increasing in important production areas of Asia since the mid-20^th^ century (Oerke 2006; Wang et al., 2022). In the last decade, Southeast Asia’s Greater Mekong Subregion (Cambodia, Lao PDR, Myanmar, Thailand, Vietnam, and the Southern Chinese provinces of Guanxi and Yunnan) has faced serious transboundary pest and pathogen invasions in major economic crops, including fall armyworm in maize (Nagoshi et al., 2020), *Fusarium oxysporum* f. sp *cubense* Tropical Race 4 in banana (Zheng et al., 2018), and cassava mosaic virus (Siriwan et al., 2020). These crops represent the livelihoods of tens of millions of smallholders across the region, and the balance sheets of major agricultural exports driving rural economies. The latter two diseases require effective seed systems to minimize disease spread.

Cassava is a staple food and industrial starch crop throughout global tropical and sub-tropical regions. In Southeast Asia, cassava is often produced by smallholder farmers on marginal or nutrient-poor land which is unsuitable for other crops (Delaquis et al., 2018; Graziosi et al., 2016), making this crop important for the livelihood of rural populations. Cassava mosaic viruses (*Geminiviridae*, *Begomovirus*), causal agents of cassava mosaic disease (CMD), are vectored by the widely distributed whitefly, *Bemisia tabaci* (*Homoptera*, *Aleyrodidae*), and dispersed in infected cassava planting stems in ineffective seed systems. Frequently recognized as one of the most important plant diseases in the tropics, the first report of CMD in Southeast Asia originated on a single cassava plantation in Cambodia in 2015 (Wang et al., 2016). Subsequent CMD spread has been reported from Vietnam (Uke et al. 2018), China (Wang et al., 2020; Wang et al., 2019), Thailand (Leiva et al. 2020), and Lao PDR (Siriwan et al. 2020).

The species of cassava mosaic virus found in Cambodia, *Sri Lankan cassava mosaic virus* (SLCMV), has previously only been reported on the Indian subcontinent, and is reported to be more virulent than *Indian cassava mosaic virus* (ICMV) (Saunders et al., 2002). Although only recently reported in Southeast Asia, cassava mosaic viruses have been a major constraint to cassava production for many years in Africa (Legg, 1999; Legg and Fauquet, 2004; Rey and Vanderschuren, 2017) and the Indian subcontinent (Saunders et al., 2002). In Africa, losses have been estimated to be greater than $1 billion (Rojas et al., 2018), with the most severe pandemic outbreak occurring in the 1990’s (Fargette et al., 2006; Legg and Fauquet, 2004). The increase in virulence during that time has been attributed to species recombination, mixed infections with more than one cassava mosaic virus species, and the occurrence of superabundant whitefly populations (Legg and Ogwal, 1998). Unlike in sub-Saharan Africa, where low-level CMD incidence had been detected for many years prior to reaching pandemic levels (Thresh and Cooter, 2005), the introduction in Southeast Asia appears to be sudden and spread has been rapid. CMD now threatens the sustainability of the regional cassava industry. The arrival of such a dangerous pathogen in a new region forces plant health authorities to make difficult decisions. How can scarce resources and personnel best be allocated for surveillance? What strategies should be adopted for the relatively small amounts of clean planting material to be impactfully disseminated in effective seed systems?

Risk-based surveillance (Cameron, 2012; Carvajal-Yepes et al., 2019; Stark et al., 2006) and management is especially critical early in a pathogen invasion, when data are limited but effective mitigation or eradication remains possible (Parnell et al., 2014). Optimizing data-driven management strategies early in an emerging outbreak is an important challenge (Epanchin-Niell et al., 2012). Early intervention is often necessary for effective eradication of an invading pathogen (Cunniffe et al., 2016), and heuristics are being developed for the best strategies for managing and surveying invasive species spreading in networks such as seed systems (Chades et al., 2011). Models have been developed for disease control in other cassava systems that are largely driven by informal planting material exchange (McQuaid et al. 2017a, McQuaid et al. 2017b).

For CMD, one of the most important management strategies is the deployment of disease-free, “clean” planting material. Due to low clonal field multiplication rates, limited volumes of vegetative planting stems can be produced by national or donor-funded programs. To maximize epidemic mitigation in the landscape, it is possible to calibrate volumes of planting material distributed, and the number and geographic location of sites to which they should be distributed. Current strategies are largely informed by local or national disease incidence, with decision making largely undertaken in the absence of comprehensive efforts to evaluate potential strategies or to visualize likely impacts of those strategies embedded in a larger regional context. There is a need for practical approaches integrating multiple elements of the disease spread cycle to provide evidence-based recommendations for seed system interventions.

Here we use a modeling framework incorporating machine learning to provide decision support for seed system management. We model environmental risk with an epidemic meta-population network model of pathogen spread through the landscape to evaluate the impact of management and surveillance campaigns. Machine learning has been used for species distribution models in ecology for many years (Drake et al., 2006; Elith and Leathwick, 2009; Lorena et al., 2011; Olden, 2008), and more recently in plant pathology to predict disease occurrence as a function of weather or climate predictors (Harteveld et al., 2017; Martinetti and Soubeyrand, 2019; Shah et al., 2014). There is great potential to predict the future geographic spread of emerging epidemics prior to the availability of complete information about pathogen epidemiology in the region (Jiménez-Valverde et al., 2011; Meentemeyer et al., 2008). Classic species distribution models are often used to model the global (or fundamental) niche of a species under the assumption that there is equilibrium in the landscape, meaning that the species has occupied all potential niches, and those that are unoccupied are not environmentally conducive for establishment. The principal problem with invasive species distribution modeling (such as must be used for emerging epidemics, like CMD in Southeast Asia) is that it is often difficult to differentiate regions where the pathogen is absent due to lack of niche suitability from those where the pathogen is absent due to lack of introduction or dispersal (Gallien et al., 2012; Jarnevich et al., 2015; Jiménez-Valverde et al., 2011; Mainali et al., 2015). Researchers have proposed solutions for this by limiting the geographic range for absence datapoints (Mainali et al., 2015; Narouei-Khandan et al., 2016), or by incorporating dispersal components (Meentemeyer et al., 2008).

In the present study, we incorporate predicted environmental risk (using machine learning) with risk of dispersal via trade in seed systems and whitefly movement. We draw on tools previously described in Andersen et al. (2019), to incorporate not only *environmental risk*, but also risk of CMD spread due to cassava stem exchange patterns and whitefly movement (*risk due to dispersal*). Network analysis has frequently been used to model the risk of pathogen spread and to identify candidate locations for sampling (Andersen Onofre et al., 2021; Buddenhagen et al., 2017; Garrett, 2021; Garrett et al., 2018; Harwood et al., 2009; Martinetti and Soubeyrand, 2019; Moslonka-Lefebvre et al., 2011; Pautasso, 2015; Pautasso and Jeger, 2008; Sanatkar et al., 2015; Shaw and Pautasso, 2014; Sutrave et al., 2012). Network analysis is particularly suited to dispersal in seed systems where nodes are geographic land units and links are the movement of planting material (and vectors) between them. For CMD in Southeast Asia, we draw on data from an empirical survey of stake exchange (Delaquis et al., 2018) to estimate network trade patterns.

We present a decision support framework that combines seed systems and environmental geospatial data layers in a multilayer network model, to forecast and map regional CMD risk in Cambodia and Vietnam, as well as in neighboring Thailand and Lao PDR. The objectives of this study are to (1) forecast disease risk in non-infected parts of Cambodia, Lao PDR, Thailand, and Vietnam by integrating disease occurence, climate, topology, and land use, using machine learning; (2) incorporate this predicted environmental risk with seed exchange survey data and whitefly spread in the landscape to model epidemic spread in a network meta-population model; and (3) use scenario analysis to identify candidate regions to be targeted for surveillance and mitigation strategies for CMD in Southeast Asia, particularly for the deployment of clean seed material.

## Methods

Our analysis had three stages, and here we provide a brief overview. First, we estimate disease risk as a function of environmental variables, comparing a set of machine learning methods. Second, we estimate a network of spread via planting material. Third, we combine these components in scenario analyses to determine the influence of management options on CMD spread. In the first component of our analysis, we use environmental predictors, coupled with CMD occurrence and absence data points from surveys conducted in Cambodia and Southern Vietnam, to predict the probability of occurrence based on environmental correlates. We draw on methods that have been developed for species distribution modeling and invasive species distribution modeling. We project risk to neighboring Thailand and Lao PDR. This portion of our analysis assumes that there are environmental differences between locations where CMD establishes and locations where it does not, and these differences are based on the ecological niche preferences of the pathogen and vector, not only lack of dispersal to these regions. The assumption that dispersal limitations do not contribute to absence may not be met completely, as we discuss more below. We compare several machine learning methods (described in detail below) to identify the best fitting model. We then incorporate this estimated environmental risk raster layer (*as a probability of establishment*), described by the best fitting model, into simulation analyses of CMD spread. Inclusion of this environmental risk parameter results in a simulation model that incorporates not only dispersal risk via trade and whitefly dispersal, but also the likelihood that the environment will be favorable for establishment.

### Invasive species distribution model of CMD establishment

#### Disease data (response variables)

SLCMV occurrence and absence data were collected from a survey conducted in 2016 in Cambodia and Vietnam (Minato et al., 2019). In brief, the virus survey was conducted in parallel with a baseline survey designed to understand farmer demographics and the regional movement of cassava planting material (Delaquis et al., 2018). In total, 419 fields were sampled from 15 districts (8 provinces) of Vietnam and 16 districts (11 provinces) of Cambodia. Districts were selected by prioritizing areas under intensive cassava cultivation. In additional to survey data from Minato et al. (2019), incidence data collected from 2016-2018 were also incorporated from the online repository pestdisplace.org associated with various CMD surveillance projects (Cuellar et al., 2018). Observations were also obtained from government surveillance campaigns in Vietnam in 2018. In total, 1022 PCR confirmed presence/absence data points were obtained from Cambodia, Vietnam, and Thailand in 2016-2018.

#### Predictor selection and dimension reduction

To model the influence of environmental predictors on CMD establishment and spread in the landscape, and to evaluate the risk of pathogen establishment in regions not yet surveyed, a series of bioclimatic, topographic, and land use predictors were assembled. Predictors were obtained from open-source repositories whenever possible. A set of 19 spatially interpolated gridded climate predictors, or climate surfaces, were obtained from WorldClim Version 2.0 (Fick and Hijmans, 2017). Additionally, monthly climate averages (temperature maximum, temperature minimum, solar radiation, average precipitation, and windspeed) were obtained from WorldClim 2.0. Elevation data were obtained from a digital elevation map (DEM) (Reuter et al., 2007). Cassava acreage data were obtained at the province or district level for Cambodia, Vietnam, Lao PDR, and Thailand from the national agricultural ministries of each country. In addition, environmental land cover estimates were obtained for 2015 from the European Space Agency (ESA) Climate Change Initiative (CCI) land cover project, from which percentage of area falling into land use categories (cropland, tree cover, shrubland, urban, etc.) were calculated. The predictors used in this analysis are described in Supplement 1.

All data were obtained as raster grids and managed in R using the packages raster (Hijmans, 2019), sf (Pebesma, 2018), rgdal (Bivand et al., 2019), and rgeos (Bivand and Rundel, 2019). All predictors were extracted from these raster layers for each of the CMD presence/absence geolocations. Models incorporating environmental variables typically run into issues of multicollinearity, which can lead to model over-fitting and poor model projection to new regions. We used the variance inflation factor (VIF) to reduce our predictor set to those that had a VIF below our defined threshold (20). This multicollinearity analysis was implemented using the vifstep function in the R package usdm (Naimi et al., 2014).

#### Classification models

Using the restricted set of predictors identified in our analysis of multicollinearity (Table 1), we compared three machine learning algorithms – random forest (RF) (Breiman, 2001), support vector machines (SVMs) (Guo et al., 2005), and XGboost (XGB) (Chen et al. 2015) – to identify the best fitting model to correctly classify the disease occurrence and absence data points. We also compared these machine learning methods to the use of logistic regression (LR).

**Table 1.**
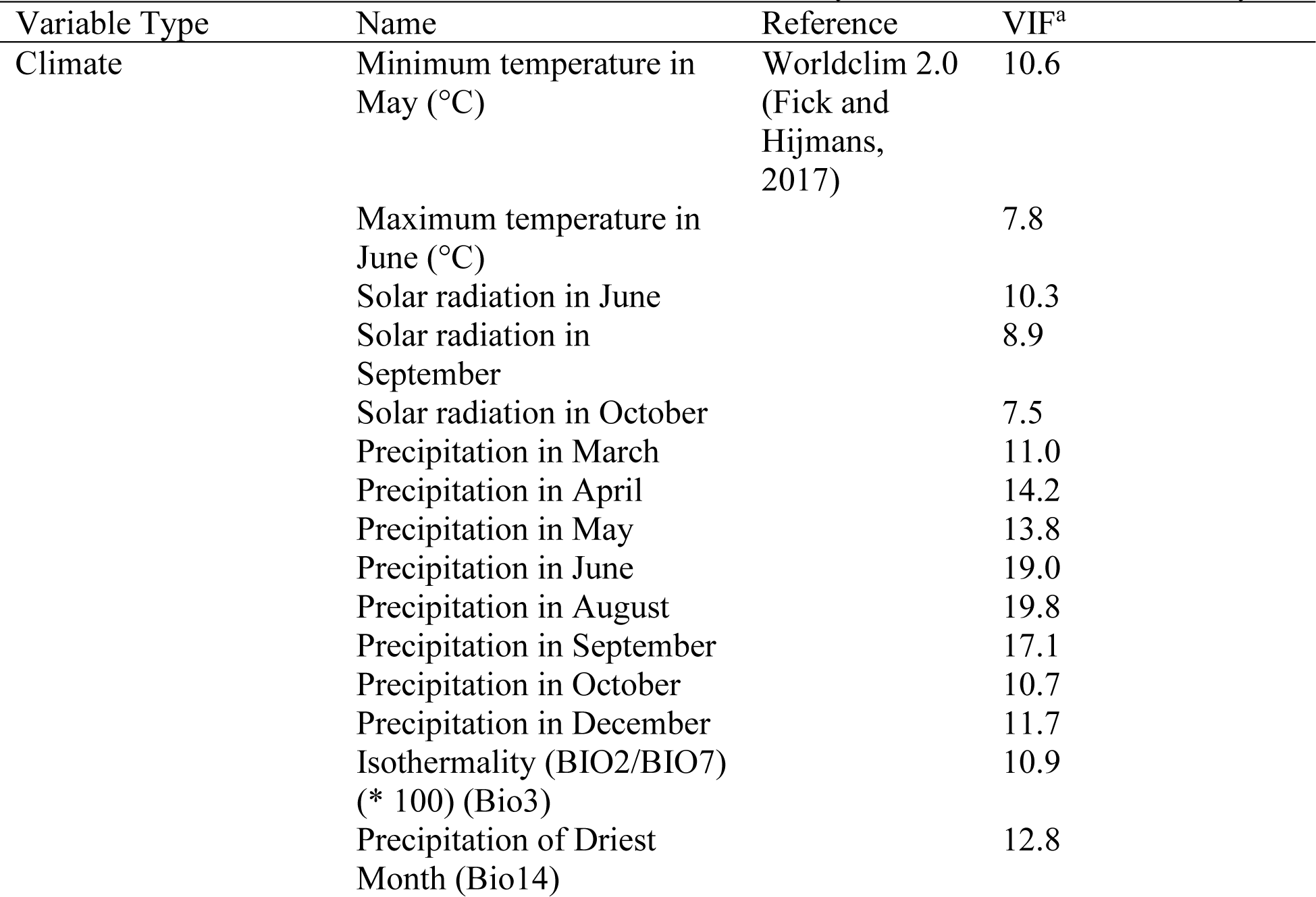
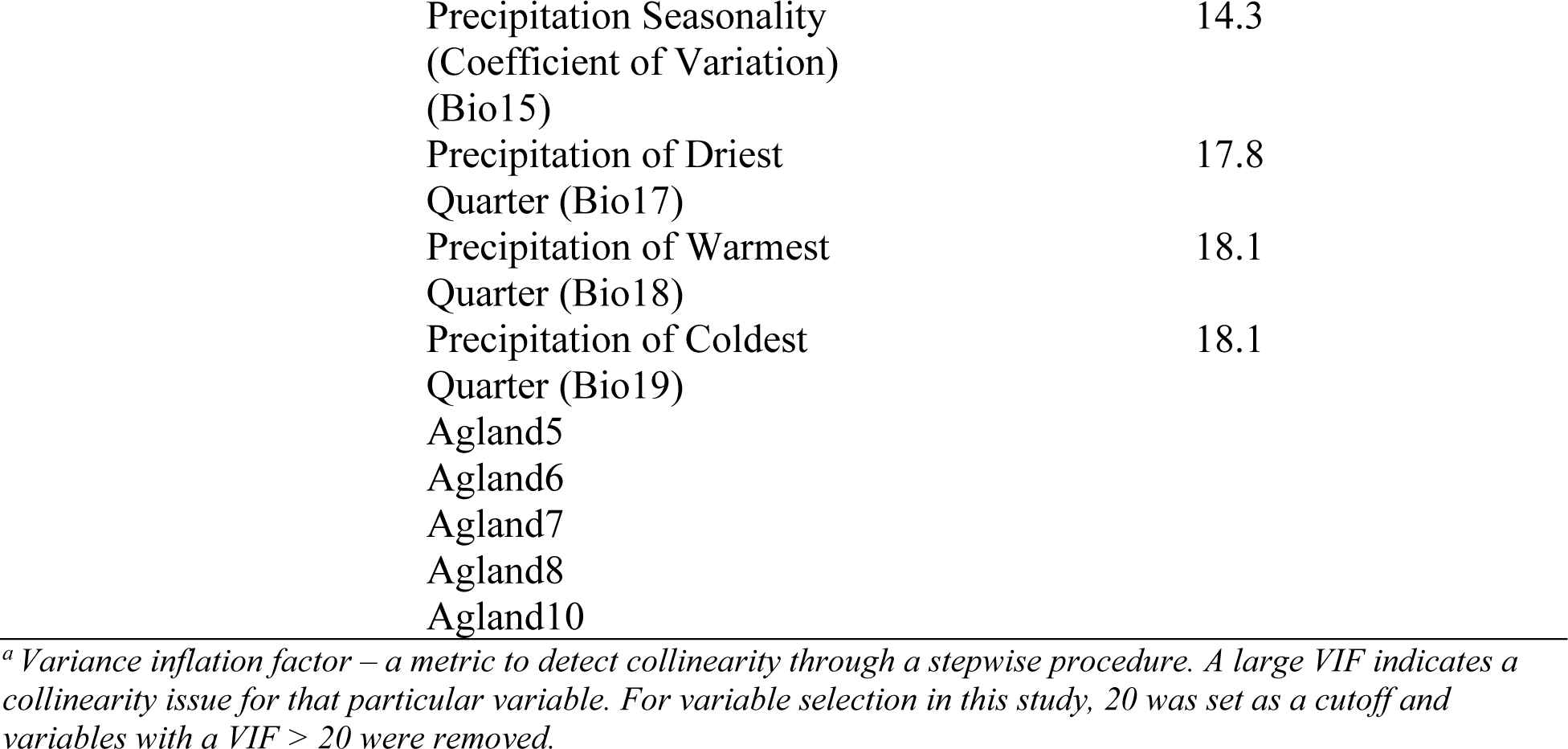
Predictors retained after variable inflation factor analysis to reduce multicollinearity.

| Variable Type | Name | Reference | VIF<sup>a</sup> |
| --- | --- | --- | --- |
| Climate | Minimum temperature in May (°C) | Worldclim 2.0 (Fick and Hijmans, 2017) | 10.6 |
|  | Maximum temperature in June (°C) |  | 7.8 |
|  | Solar radiation in June |  | 10.3 |
|  | Solar radiation in September |  | 8.9 |
|  | Solar radiation in October |  | 7.5 |
|  | Precipitation in March |  | 11.0 |
|  | Precipitation in April |  | 14.2 |
|  | Precipitation in May |  | 13.8 |
|  | Precipitation in June |  | 19.0 |
|  | Precipitation in August |  | 19.8 |
|  | Precipitation in September |  | 17.1 |
|  | Precipitation in October |  | 10.7 |
|  | Precipitation in December |  | 11.7 |
|  | Isothermality (BIO2/BIO7) (* 100) (Bio3) |  | 10.9 |
|  | Precipitation of Driest Month (Bio14) |  | 12.8 |

|  |  |
| --- | --- |
| Precipitation Seasonality<br>(Coefficient of Variation)<br>(Bio15) | 14.3 |
| Precipitation of Driest<br>Quarter (Bio17) | 17.8 |
| Precipitation of Warmest<br>Quarter (Bio18) | 18.1 |
| Precipitation of Coldest<br>Quarter (Bio19) | 18.1 |
| Agland5 |  |
| Agland6 |  |
| Agland7 |  |
| Agland8 |  |
| Agland10 |  |

<sup>a</sup> Variance inflation factor – a metric to detect collinearity through a stepwise procedure. A large VIF indicates a collinearity issue for that particular variable. For variable selection in this study, 20 was set as a cutoff and variables with a VIF > 20 were removed.

#### Machine learning and GAMs

RF, SVMs, and LR were implemented in the mlr (Bischl et al. 2016) package in R. All models were evaluated through k-fold cross validation (10-fold), repeated 50 times. The model was evaluated using average AUC, TSS, and Cohen’s Kappa statistics, common metrics for evaluating ecological species distribution model accuracy (Allouche et al., 2006) (Table 2).

**Table 2.** Classification model performance.

| Method | AUC | TSS | Cohen's Kappa |
| --- | --- | --- | --- |
| Random Forest | 0.94 | 0.52 | 0.62 |
| Logistic regression | 0.80 | 0.22 | 0.30 |
| SVMs | 0.83 | 0.38 | 0.45 |

#### CMD risk projection

Once the best model was established (based on AUC, TSS, and Cohen’s Kappa), it was used to project predicted probability of establishment to the remainder of the region. Predictions were made based on a random gridded sample of the uninfected region, 3000 geolocations in total, throughout unsampled regions of Cambodia and Vietnam, as well as Thailand and Lao PDR, two important neighboring regions with growing cassava production. This average probability per district was evaluated and is denoted as risk of establishment, ε. Additionally, a scaling parameter, *α*, was added to simulation models (described below) to modify the influence of the environment on establishment. Districts where no cassava acreage was reported in the previous three years were not included in this analysis.

### Network model of CMD spread

Environmental conduciveness (described above) is only one component of the risk of CMD spread. Dispersal, through the movement of virus-infected planting material in seed systems and via the movement of viruliferous whiteflies, also plays an important role in determining where CMD is present in the landscape. We estimate the likelihood of both modes of dispersal, as described below, and combine them for use in our simulation experiments. Once the simulation model was constructed, several scenarios for clean seed deployment were tested. The epidemic model that we implement is similar to that in Andersen et al. (2019), using a discrete time network SI (*susceptible-infected*) Markov chain, with the following changes to adapt the model to this system. First, we incorporate within-node disease spread dynamics as a function of transmission (rate β) between healthy and infected hectarage within a district (node) and between neighboring districts, representing local spread via whiteflies (Figure 1). Second, the probability of infection is not based solely on the probability of transmission, but also on the probability of establishment, ε (described above). Finally, in a management/intervention scenario analysis, we introduce incomplete recovery at nodes, as a function of the proportion healthy cassava planting material introduced into select nodes at the start of the season.

**Figure 1.**
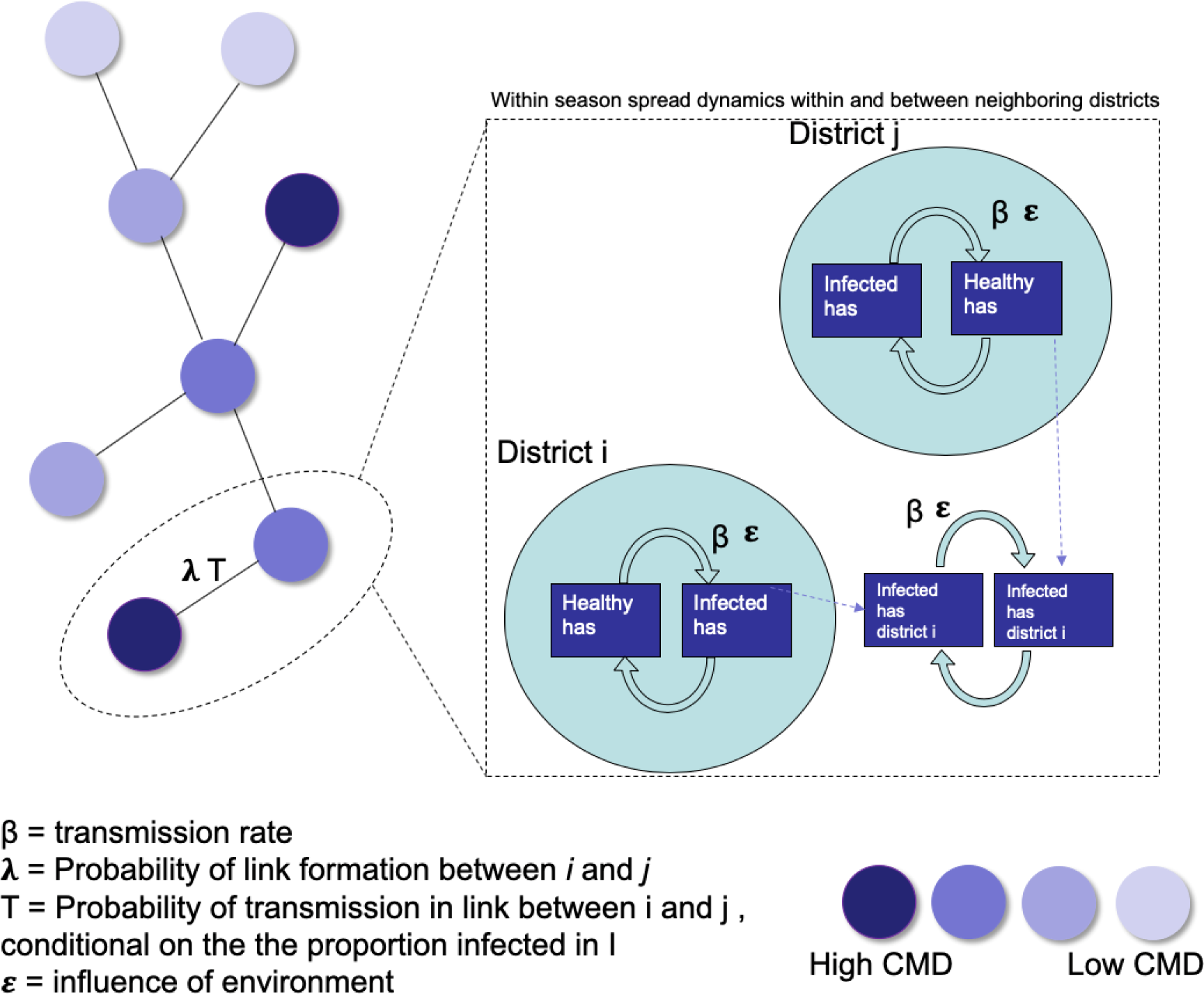
Network metapopulation model schematic for cassava mosaic disease in seed systems and via vectors. The network illustration on the left side of the figure represents cassava seed trade links at the start of the season. If cassava planting material is moved from an infected district, new cassava areas can become infected in the “sink” district. The box on the right side of the figure illustrates within- and between-node spread dynamics during the course of the season. Infected planting material can infect healthy cassava areas within a district, but can also infect healthy cassava areas in adjacent districts. Within-season spread represents whitefly transmission. Between-node links (representing cassava seed trade) only occur at the initiation of a season. Within-node spread, depicted in the breakout box, occurs over the course of 10 monthly timesteps within the season.

### Model Components

#### Seed system network

We fit a network model of planting stem exchange in the region (Cambodia, Vietnam, Lao PDR, and Thailand) where nodes are aggregated to the scale of districts (sub-provincial administrative units) and links are directed and calculated based on the probability of stem exchange between nodes (districts). In a related study, data were collected on farmer behavior related to planting material exchange (Delaquis et al., 2018) and these data were used to estimate link probabilities. The probability of a link forming between any two districts *i* and *j* (λ_ij_) was estimated based on the household survey results. The frequency with which a link was reported between two districts for each of a set of distance ranges (0-100, 100-300, and 300-500 km) was calculated, and then the product of this frequency and the total number of districts at that distance apart was taken as the probability that a link would form in each season. This incorporates a low likelihood that a district would trade with *each* neighbor, but a district would tend to trade with *some* neighbors in each season. No trade events were observed at > 500 km distance, so probabilities of exchange between nodes > 500 km apart were set to zero.

#### Short-distance whitefly dispersal

We are not only concerned with CMD spread in the landscape via seed systems, but also via the movement of viruliferous whiteflies. It has been documented that the bulk of whiteflies disperse anisotropically in the landscape up to a maximum of 2 km from their source (Byrne et al., 1996), with a majority not migrating from their source. Because the nodes in our analysis represent districts, we consider whitefly movement to be important only between adjacent districts within the month-long timesteps of the model, and thus only make it possible for whiteflies to move between adjacent districts within timesteps. In the next timestep, new generations of whiteflies may emerge and infect adjacent districts. Unlike cassava planting stem exchange, which only occurs at the start of a season, whitefly spread can occur throughout the course of a season. We account for this by allowing movement of whiteflies between districts to occur over a monthly time step, with 10 timesteps during the season corresponding to a roughly 10-month cassava growing season. In this scenario, whiteflies from adjacent districts may move into neighboring districts and cause infection of previously healthy material there. In the next month there is the potential for these whiteflies to advance from the newly infected district to the next neighboring district. This process continues until the end of the season, at which time the final amount of infected area is maintained to restart the next season.

### Simulation model description

In our simulation model, each node (district within Vietnam, Cambodia, Lao PDR, and Thailand which reported cassava planted area in 2018, 1255 in total) begins with a fixed number of hectares of cassava, corresponding to the hectares of planted cassava reported for the district in 2018 (CIAT 2018). Additional starting conditions for the simulation include CMD infection status of each district, hectares infected with CMD (*D*_*it*_), and corresponding hectares uninfected (*H*_*it*_). All districts reported to have CMD infection (based on previous survey results, Ministry of Agriculture reports, and the opinion of experts from the region) were designated as infected. In areas where CMD was only reported at the scale of the province, three districts with the highest number of planted hectares were designated as CMD positive. In total, 125 districts were CMD positive in the start of the simulations, reflecting the estimated CMD prevalence at the close of the 2019 season. Districts were each assigned a 1% incidence to begin (1% of hectares infected). This is likely a conservative estimate of total infected hectares at the close of the 2019 season.

We describe the infection process in node *i* by which a portion of previously healthy hectares of cassava becomes infected during a month-long timestep to yield a newly infected total number of hectares with CMD (*D*_*i*,(t+1)_) as

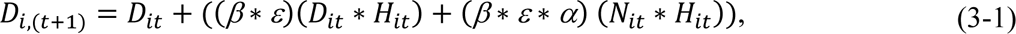

where *D*_*it*_ is the number of infected hectares in a district in district *i* in month *t*, *H*_*it*_ is the number of healthy hectares in district *i* in month *t*, *H*_*it*_ is the sum of infected hectares in all districts adjacent to district *i* (external inoculum sources), *β* is the transmission rate of infection from infected to uninfected hectares via whitefly spread, *ε* is a parameter describing the environmental conduciveness to disease (based on the species distribution modeling), and *α* is the relative influence of neighboring inoculum compared to within-district inoculum. The healthy area in the next timestep (month) was

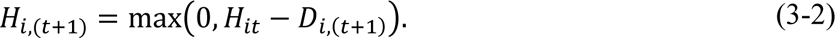

At the close of each timestep, *H*_*it*_ is the sum of infected hectares in districts adjacent to district *i*

(external inoculum sources), calculated as

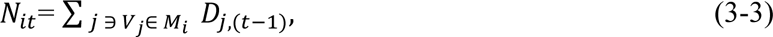

where *V_j_* is node *j* and *M_i_* is the set of nodes linked to node *i*. This simulation is carried out over the course of 10 timesteps, corresponding to the 10-month typical cassava growing season in this region.

At the end of each season *t*, a set of ending conditions for each district was obtained. At this point a network model of the seed system was implemented, simulating the seed exchange that takes place prior to planting at the start of a new season (*t* + 1). Exchange of planting material occurs across network trade links (where *λ_ij_*, the probability of trade between districts *i* and *j*, is used to stochastically generate trade events as described above). If a trade event occurs from node *i* (the source node) to node *j*, the likelihood that the material moved is infected with CMD was set to 10%. This infected hectarage that is “moved” to sink node *j* (set to 5 hectares) is then summed with the infected hectares of node *j* (*D*_*it*_) for the start of the new season (and the corresponding *H*_it_ is also updated). Five hectares was chosen as the trade volume based on previous survey results where the average amount of stems per transaction was 11,000 (Delaquis et al., 2018). On average it takes approximately 2,000 stems to plant a hectare of cassava in this region, so five hectares was determined to be a reasonable area to be introduced by trade. For this subsequent season, after seed exchange, a new round of within season simulations is started, as described above. At the end of the season *t*, a new *D*_*i*,(t+1)_ and *H*_*i*,(t+1)_ are stored for each node, and are used as the starting conditions for the next season, in which the process is repeated. Simulations were run over 10 seasons with 1000 realizations per parameter combination. Total number of hectares infected in the region, along with total number of districts infected, was evaluated.

### Uncertainty

A sensitivity analysis (uncertainty quantification) was conducted for two of the model parameters, *α* and *β*. The transmission parameter, *β*, represents the rate at which uninfected hectares become infected by within-district infected and neighboring infected hectares, reflecting the spread of CMD via infected whiteflies. Because insufficient data was available to estimate this parameter, a range of values was tested (0.0002, 0.00002, 0.000002, 0.0000002).

Additionally, a range of values of the parameter *α* (0.1, 0.3, 0.5, 0.7,0.9), which modifies the effect of the inoculum of neighboring districts in the model, was tested. Model output was evaluated for each pairwise combination of these parameter values.

### Simulation Experiments

Using the above-described network model, we carried out simulation experiments with corresponding sensitivity analyses to address the following key questions: 1) How does completely restricting stem trade influence the spread of CMD in the region, 2) how does incorporating clean seed into the system each year impact trade, as a function of clean seed volume, and 3) how should locations for clean seed interventions be selected?

#### Experiment one: seed trade restriction

A scenario with no stem exchange was evaluated. In this “trade restricted” simulation, planting material that was infected at the end of the previous season (t-1) was used as the starting amount of planting material for the current season (t), over the course of ten seasons. Like all scenarios in this study, we assumed no positive selection or reversion and thus no reduction in infected area from year to year. This set of simulations was deterministic (because without trade, the stochastic components of the model are not engaged). The percent reduction in infected area across all districts and total number of infected districts was calculated by comparing the outcomes from the simulations described above for “full trade” with these “trade restricted” scenarios.

#### Experiment two: clean seed intervention strategies

Clean seed is one of the most effective ways to manage CMD and other seed transmitted diseases of vegetatively propagated crops. Plants from clean planting stems that become infected with CMD later in the season lose less yield than plants from infected planting material (Fauquet and Fargette, 1990). However, the capacity for clean stem production is a limiting factor, thus an urgent and practical question is: What are the optimal locations, and corresponding volumes, for clean cassava planting material deployment? In this region, it typically takes 10,000 cuttings to replant one cassava hectare (where five cuttings are obtained per stem for most varieties). In 2019, Vietnam’s government reported over 31,000 ha infected by CMD with varying levels of severity, requiring >300,000,000 cuttings to replace this area with clean planting material. Producing this volume of clean seed material is currently not feasible, so there is a need to prioritize deployment in the region, to maximize effect.

We introduce “management” into the system via the infusion of “clean planting material”. This material, in the form of stems, is assumed to be certified disease free, from stock originating in tissue culture and bulked in the field where there was no risk of infection. After the exchange event in the model, as part of the start of the season (*t*), a set volume of added clean planting material is summed with the clean material already present in a district to be managed, and removed from the proportion of infected hectares *D*_*i*,(t+1)_for the district from the end of the previous season. Here we make several assumptions: 1) that the proportion of infected planting material from the previous year is otherwise retained, except for what is added as clean planting material, 2) that the amount of infected hectares at the end of the previous season is equal to the proportion of infected hectares at the start of the next season (farmers do not change their cassava acreage from season to season, and farmers do not employ positive or negative selection on saved planting material), 3) clean stems will completely replace a proportion of infected hectares at the start of the season, 4) cassava from clean stems can become infected during the season at the same rate as all other cassava.

We test three methods for selecting locations to manage: 1) “random selection in infection zones” where cassava-producing districts that fall in infection zones (infected themselves and/or within a 50 km buffer of infected districts) are randomly selected in each season, 2) districts are selected within the same infection zones as described in scenario one based on their starting area of diseased material (those with highest area selected first), and are managed each season, and 3) locations are selected based on a reactive management rule – each season locations are prioritized based on their infected area in the previous season, and again areas with the most infection are managed first. We not only wanted to understand the best strategy for managing locations, but also the optimal number of locations and volume to distribute to each location. We tested managing 20, 40, 60, and 80 districts, with each combination of 20, 40, 60 and 80 ha of clean planting stems. For example, for one treatment combination, 20 locations would be managed with 20 ha each of clean planting stems, and so on. Each management scenario (combination of (a) method of selecting locations, (b) number of locations, and (c) volume of disease-free stems) was evaluated, and scenarios were compared for their utility in slowing the spread of CMD through the region over time. We implemented these management scenarios for the two most conservative parameter estimates: *β* = 0.000002 and *α* = 0.1.

The code for these analyses will be available in GitHub upon publication. Key aspects of the analyses are being incorporated as part of the R2M toolbox (www.garrettlab.com/r2m), which addresses rapid risk assessment to support mitigation of plant pathogens and pests (Andersen et al., 2019; Andersen Onofre et al., 2021; Buddenhagen et al., 2017; Buddenhagen et al., 2022; Etherton et al., 2023; Garrett, 2021; Margosian et al., 2009; Nduwimana et al., 2022; Xing et al., 2020).

### Model feedback and stakeholder participation

Initial model assumptions, underlying data, and preliminary model outputs were described to model stakeholders during a session of a workshop held in Vientiane, Lao PDR on September 12^th^, 2019. In addition to CIAT researchers and the study authors, there were participants from local universities, private cassava product companies, and national plant protection programs. The workshop feedback survey is in Supplement 2.

## Results

### Environmental predictor selection and model performance

From the 98 input variables that were initially considered, 78 were removed prior to modeling due to multicollinearity. The remaining 24 variables were included in downstream analyses (Table 1). Among the models tested, random forest had the highest AUC, TSS and Kappa values and was used for calculating district-wide mean establishment risk estimates (*ε*) (Table 2). The best-fitting random forest model was used to project risk to the remainder of the region. Average district-level establishment risk varied from 0.05 to 0.94 (Figure 2).

**Figure 2.**
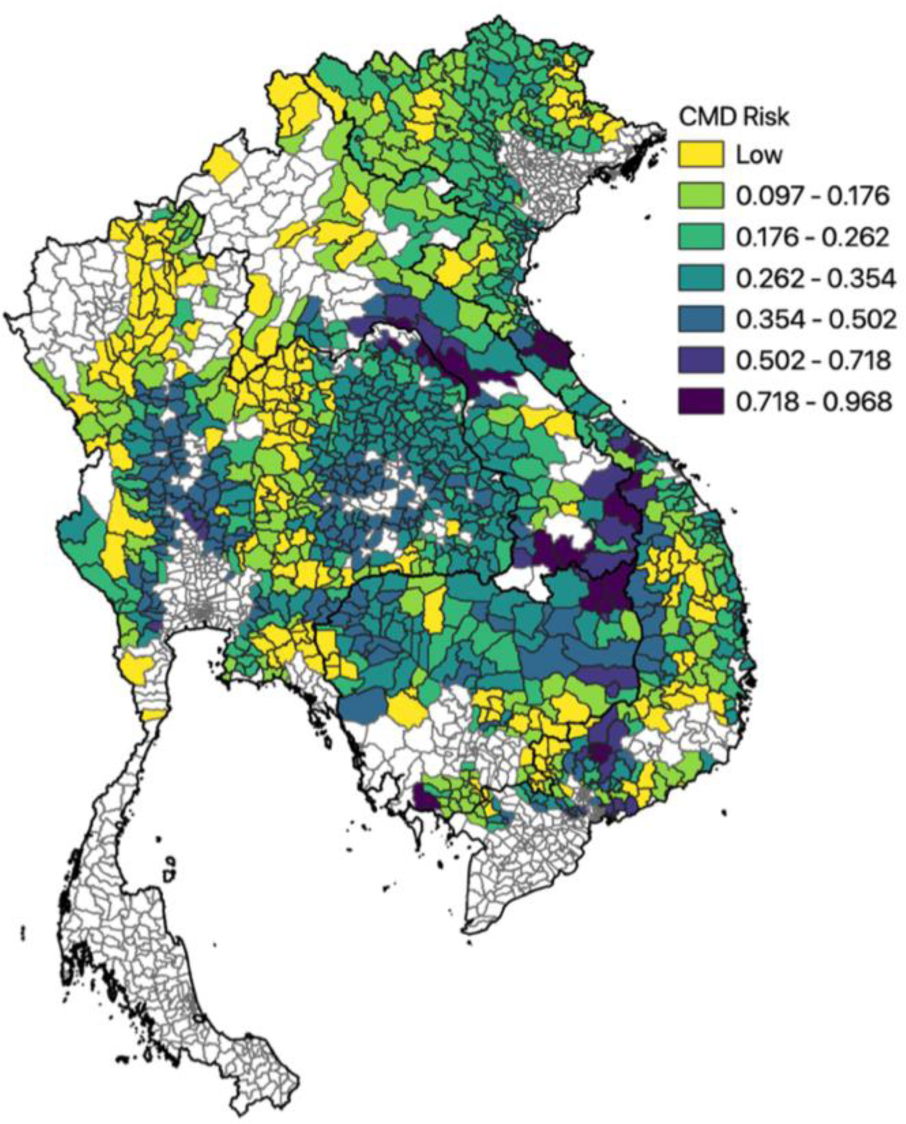
Estimated cassava mosaic disease (CMD) risk in SE Asia. District estimates are based on the average of approximately five point estimates selected randomly from within a district. Low scores (light color) represent low CMD risk, and high scores (dark colors), represent high risk.

### Epidemic spread and model parameter sensitivity

Simulations of CMD spread, with 125 districts starting as infected, were conducted with all pairwise combinations of *β* and *α* (Figure 3). Lower values of *β* give lower rates of disease spread. For example, in the scenario of the lowest *α* value, 0.1, and lowest *β* value, 0.000002, after the completion of the first season a mean (across simulations) of 419 districts had some level of CMD incidence (min = 417, max = 431). By season 5, the mean was 489 districts infected (min = 482, max = 512). The total area infected for the same parameter combination was a mean of 7,679 ha (min = 7674 ha, max = 7695 ha) with CMD, which is approximately 0.2% of total cassava area in these four countries. By season 5, the mean number of hectares infected was 10,253 (min = 10255, max = 10304). In contrast, for the scenarios with the highest values of *β* and *α*, 0.002 and 0.9, respectively, the mean number of districts infected by the close of the first season is 922 (min = 920, max = 975), or 73% of all districts across the four countries. At the end of this first season, over 2 million hectares of cassava in the region have some level of infection, or 82% of cassava producing area, and this number jumps to 2.4 million by the end of the 5^th^ season.

**Figure 3.**
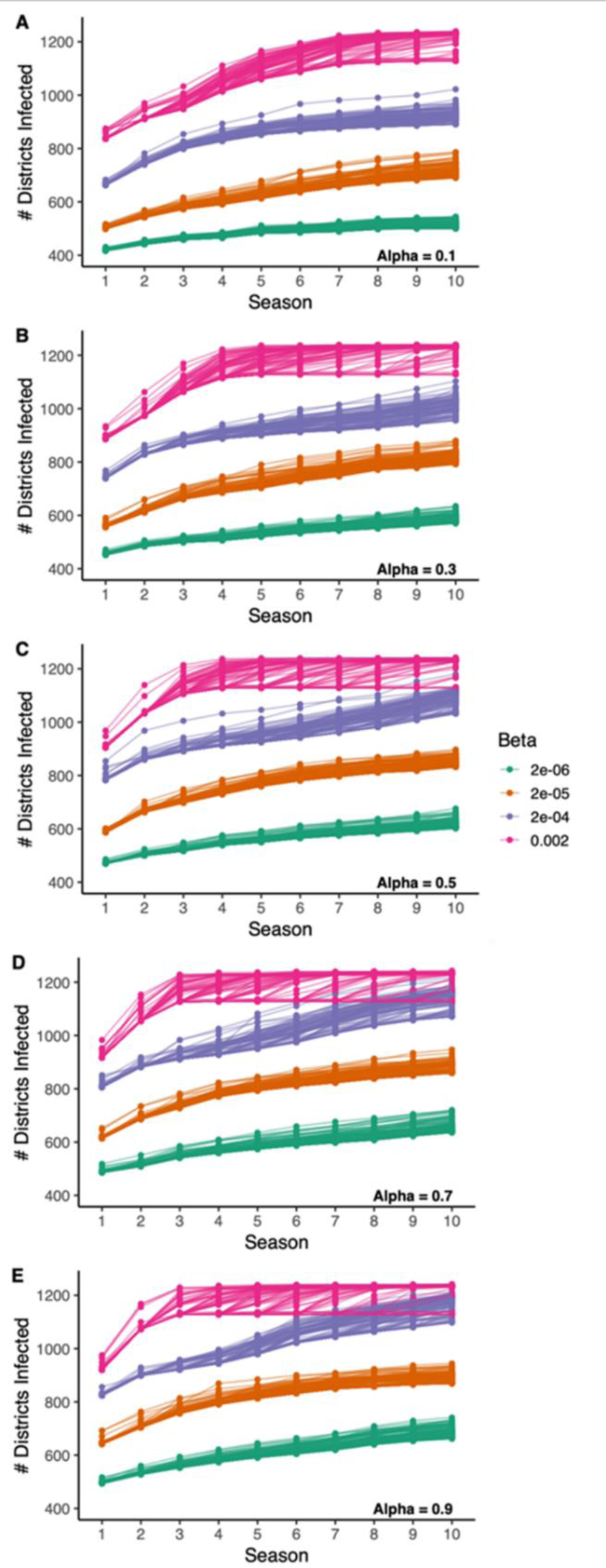
Cassava mosaic disease (CMD) in Southeast Asia: pathogen spread in simulations over 10 seasons for each combinations of four values of a dispersal parameter (*β*) and five values of a parameter (*α*) modifying the effect of the environment (ε) and external inoculum. Points indicate the number of districts with CMD present in 100 realizations.

### Effect of halting seed trade

In an initial scenario analysis, to assess the upper limits of management efficacy in the system, we addressed the question of how much a complete trade restriction would limit pathogen spread. In this scenario, the only potential for spread was via spread of viruliferous whiteflies in the landscape. As could be expected, this slowed the epidemic for all parameter combinations, when compared to scenarios of free trade (Figure 4). Across all the parameter values, the percent reduction in infected hectares ranged from 4% to 92% with an average reduction of 46%, across all parameter combinations and timesteps.

**Figure 4.**
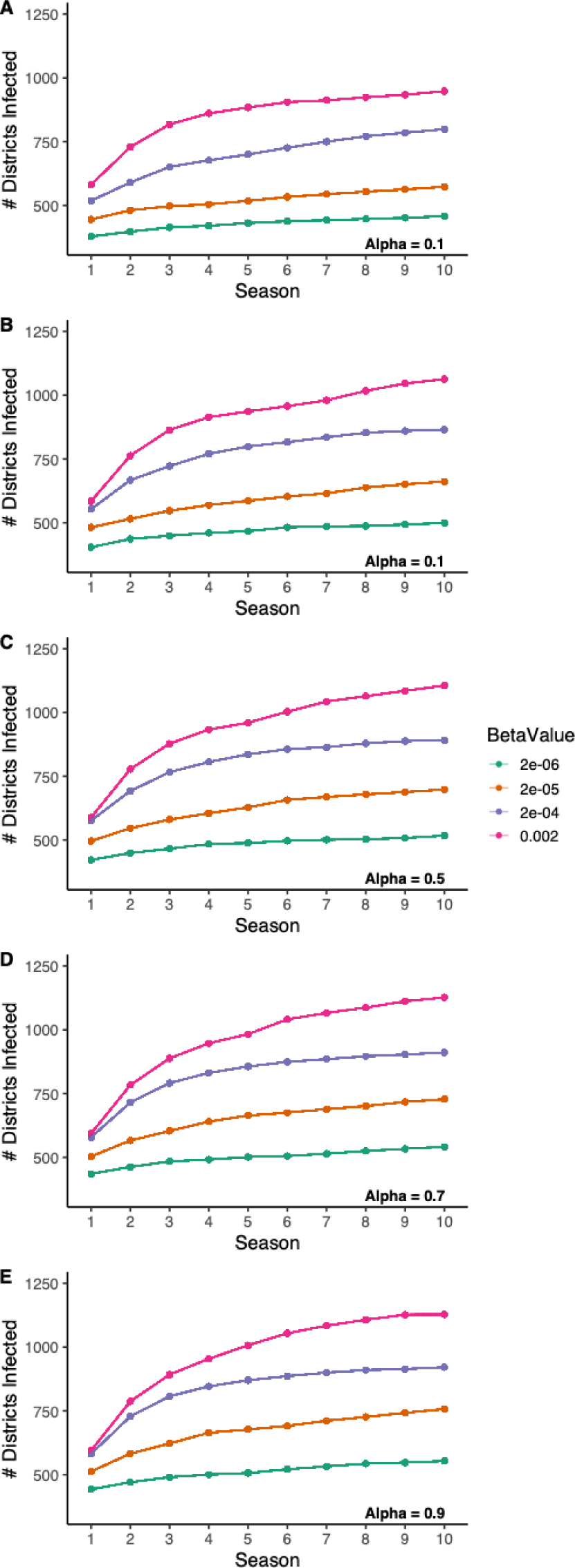
Cassava mosaic disease (CMD) in Southeast Asia **with seed trade halted** so that spread is solely due to vectors. Pathogen spread is simulated over 10 seasons for each combinations of four values of a dispersal parameter (*β*) and five values of a parameter (*α*) modifying the effect of the environment (ε) and external inoculum. Points indicate the number of districts with CMD present in 100 realizations.

### Clean Seed Provisioning Strategies

We tested three scenarios for selecting locations for clean seed deployment, as well as all combinations of managing 20, 40, 60, and 80 districts and 20, 40, 60, and 80 hectares. The least effective method for selecting locations to manage was scenario one, randomly selecting districts within 50 km of infected areas without consideration of infection status (Figure 5a). Although in some realizations this method did perform well, the results were highly variable across realizations. This high variability was likely due to new locations being selected in each realization, which may or may not have been districts with previous CMD infection.

**Figure 5a-c.**
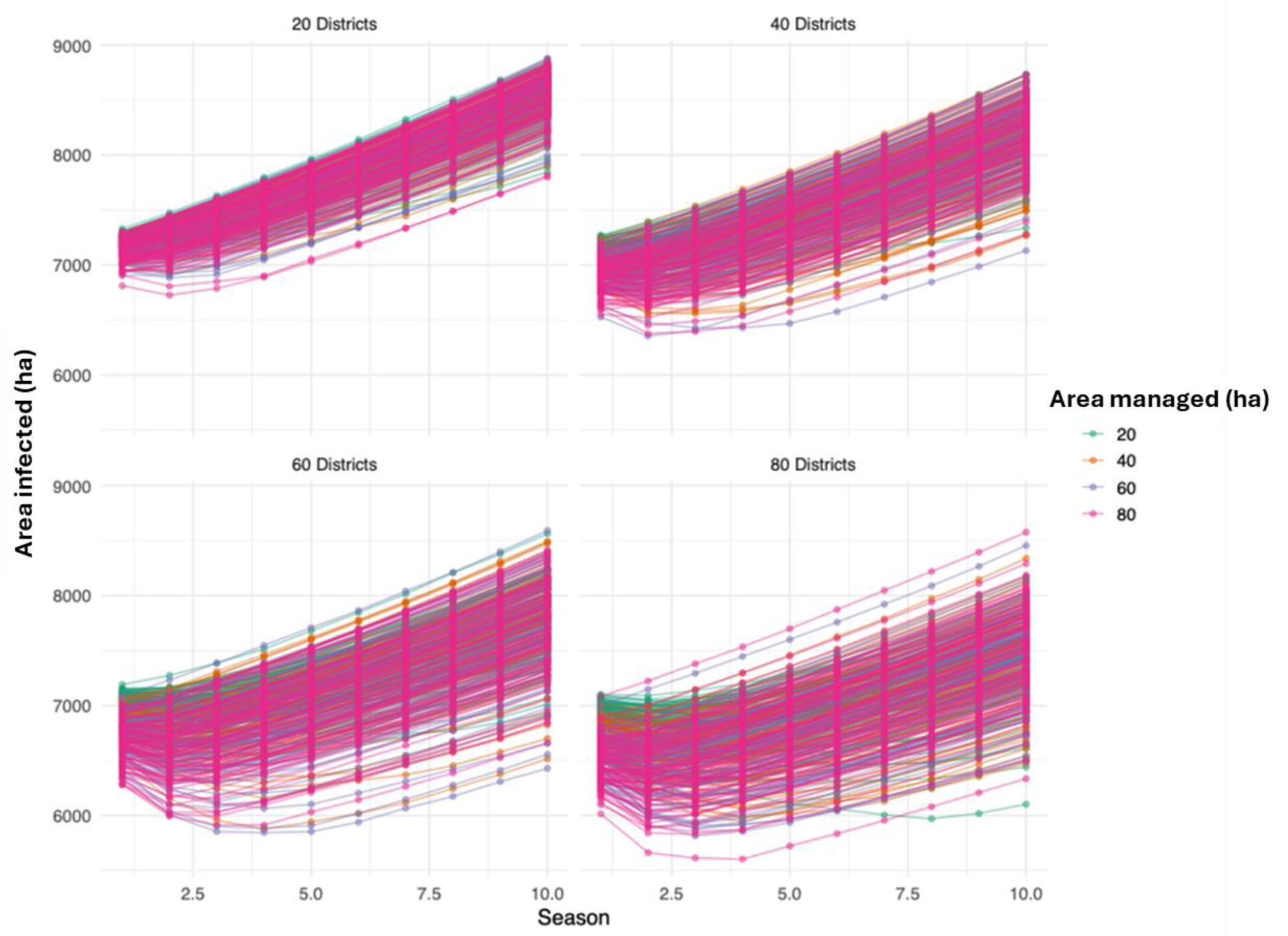
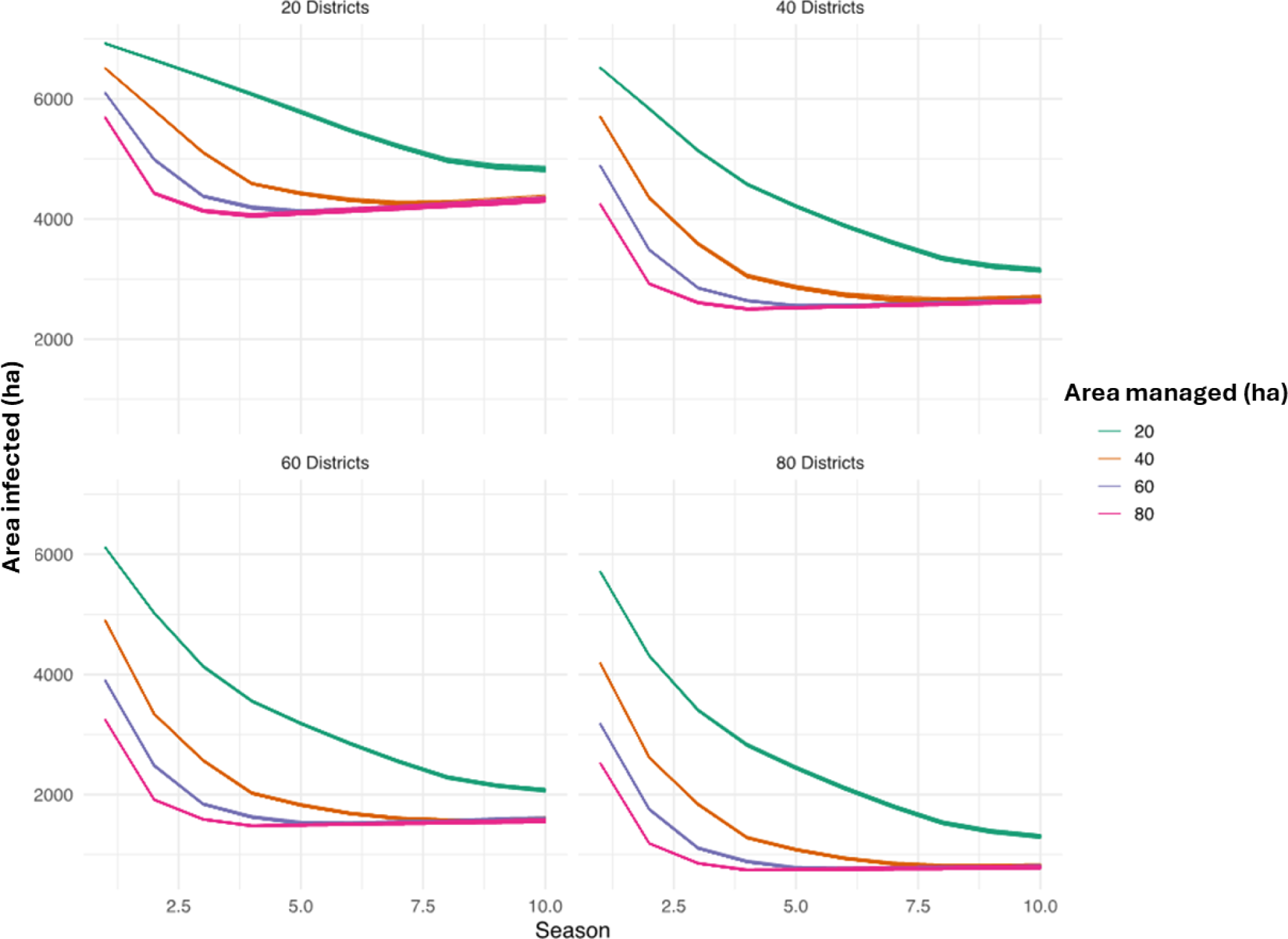
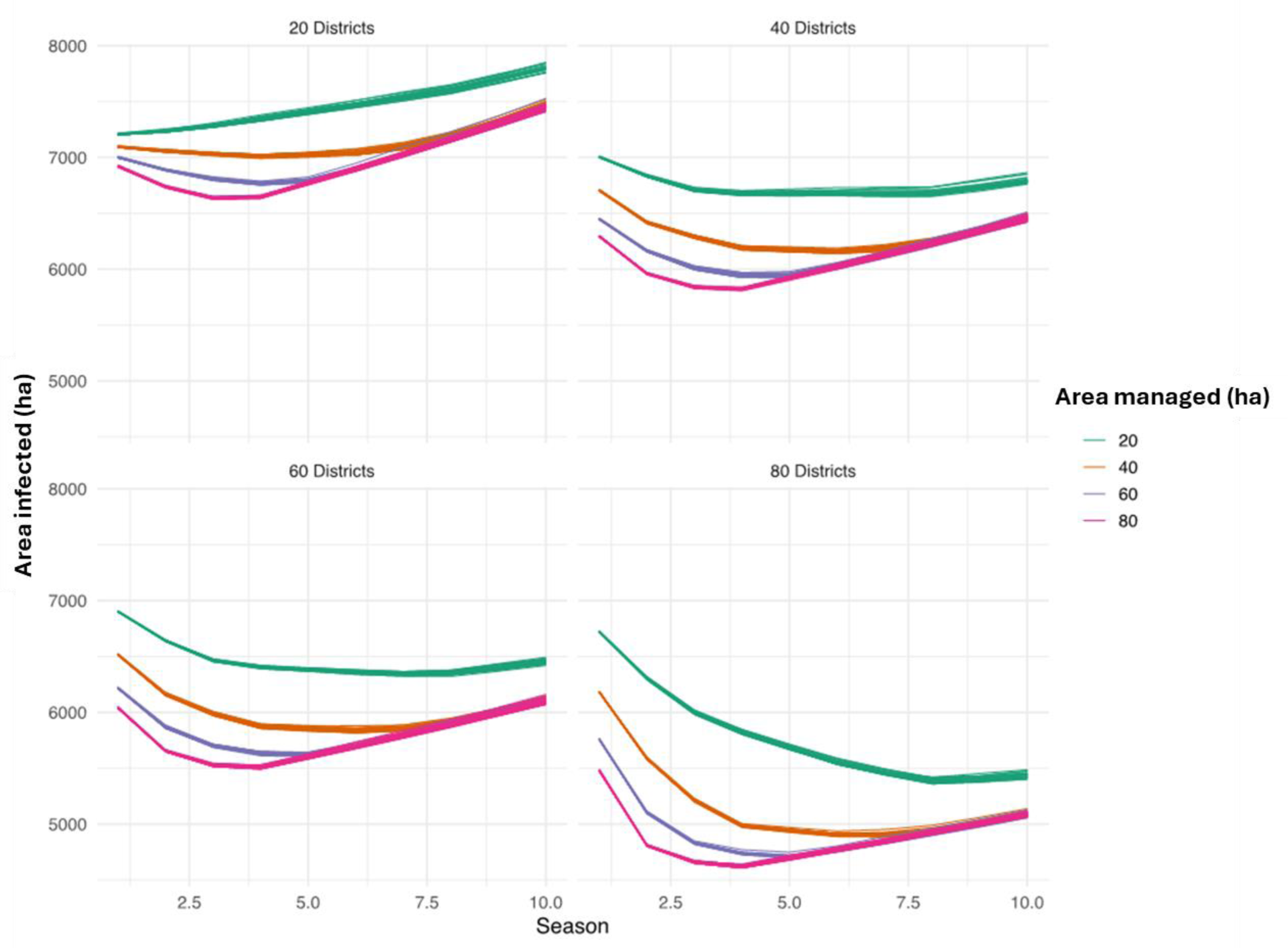
Cassava mosaic disease (CMD) in Southeast Asia for simulations of pathogen spread **in three scenarios of clean seed provisioning**. A) In the first scenario, with “random management” clean seed is deployed in randomly selected locations within a 50km radius of infected districts. B) In the second scenario, the selection of locations for clean seed deployment was based on the initial area with CMD, and selected locations were managed each season. C) In the third scenario. “reactive management”, locations with the greatest area with CMD were selected independently each year, with the potential to change across time. Individual plots give the simulation outcomes for sub-scenarios in which 20, 40, 60 or 80 districts are managed, with the number of ha managed per district indicated by color.

The best strategy for reducing CMD was scenario two, selecting locations at the start of the season with the highest number of infected hectares to manage and managing them each season with the same volume of clean planting material, a “fixed location campaign” (Figure 5b). This scenario also has important practical benefits, as it is easier to manage a fixed number of highly infected areas in a concentrated campaign compared to constantly seeking out new regions to manage. Fixed locations are also more logistically realistic given the stationary nature of most planting material multiplication infrastructure, paired with the relatively high transport cost of bulky cassava stems. Interestingly, the fixed location campaign was more effective than the “reactive management campaign”, where locations for management were selected at the start of each season based on the number of locations that had the highest number of infected hectares in the previous season. The reactive strategy results in shifting locations to manage, allowing for more hectares of infected area in some cases. For the “fixed location campaign” and the “reactive management campaign” the variation from simulation to simulation was relatively low (Figure 5a and Figure 5b). This is in part due to the relatively small infected area and low rate of increase due to the choice of the conservative parameter estimates *α* = .1 and *β* = 0.000002.

An important component of this analysis was identifying the best strategy for prioritizing locations to manage, and total area to manage, given a limited amount of available planting material. For our “fixed location campaign” managing only 20 locations, even 20 ha led to a decline in CMD prevalence season after season (Figure 5b), with a more rapid decline occurring when 40, 60, or 80 ha were managed for each of the 20 locations. As the number of districts managed increased, the decline in CMD prevalence was more pronounced. Interestingly, in all cases, managing 40, 60, and 80 hectares gave similar results, suggesting that an intermediate volume of clean planting material might be sufficient to lower spread.

### Stakeholder workshop feedback

Generally, stakeholders were positive about the utility of the model results (Figure 6). Among survey respondents, 75-80% indicated that model results would be useful for planning surveillance and mitigation campaigns. Most agreed that the model results were easy to interpret. Several useful suggestions were left in open-ended comments on the worksheet. This stakeholder feedback was carefully considered in subsequent rounds of analysis and revision of this work.

**Figure 6.**
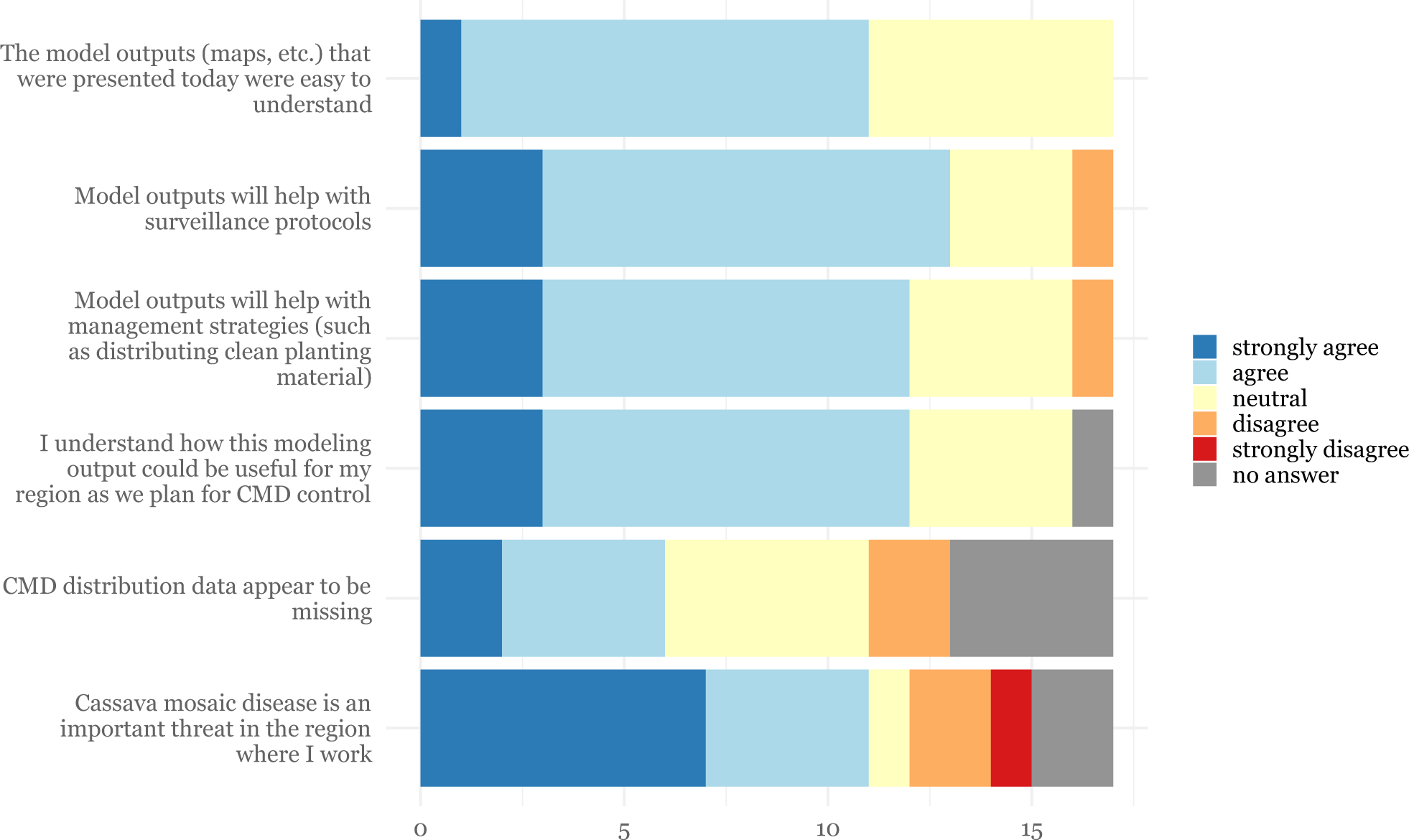
Workshop stakeholder feedback about the structure and results from a model of cassava mosaic disease in Southeast Asia designed to support decision making about clean seed deployment to limit pathogen spread. A feedback session was conducted during a workshop with stakeholders from Vietnam, Cambodia, Lao PDR, Thailand, and China. Relative numbers of responses in each category for each question are shown. There were 35 participants, and 17 survey responses.

## Discussion

Regional deployment of clean seed is an important management component for emerging seedborne pathogens. This study supports decision-making about how to prioritize locations for clean seed deployment. In the simuliations, consistently managing even a small percentage of the 1255 cassava-producing districts in the Greater Mekong subregion, when selected based on being among those with the highest prevalence of CMD, could reduce disease year after year and result in an overall slowing of spread in the sub-region. Even minor reductions in the rate of spread in the region can allow for additional time to prepare control measures and conduct research. The consistent management of highly infected districts greatly outperformed consistent management of randomly selected districts near infected areas. Managing highly infected areas consistently also outperformed the ‘reactive management’ model in which locations changed based on seasonal disease levels.

The underperformance of the reactive management regime may be due to the incomplete disease eradication resulting from limited resources, allowing target districts to become re-infected in subsequent seasons. Additionally, a small fraction of districts (maximum 80) were managed at any given time, which is ∼6% of the total number of districts in the system. Once the epidemic escapes from the initially infected area, management becomes less effective (accounting for the shift from a decrease in total area infected to an increase). It should be noted that districts to manage were selected solely based on the area infected, not accounting for spatial autocorrelation of districts. Selecting districts to manage by taking into account their spatial proximity (for example, managing adjacent districts or “foci”) might improve the method, and would be a good topic for future work.

Informal seed systems for crops that are vegetatively propagated, like cassava, can move viruses rapidly. In Southeast Asian, cassava stakes commonly move through informal seed systems, without regard for international boundaries, and often over distances > 400 km (Delaquis et al., 2018). The model scenarios of trade restriction showed a dramatic reduction in CMD spread when trade was restricted so that movement was limited to local whitefly dispersal and transmission in the landscape. Trade restrictions could dramatically reduce spread, particularly to regions not adjacent to infected areas. Although informal trade is legally restricted in several Greater Mekong Subregion countries, laws are difficult or impossible to enforce completely in clonally propagated crops, and informal seed systems remain the backbone of the cassava sector. Thus, seed trade policies alone are not sufficient to control CMD in the region.

The invasive species distribution model of CMD found climate and land use to be suitable predictors for estimating the potential distribution range in the region. In this study we compared the efficacy of several machine learning techniques that are generally well suited for classification of invasive species distribution data, with the ability to handle a large suite of explanatory predictors: the maximum entropy model (Maxent), random forest (RF), and support vector machines (SVMs). Maxent (Phillips et al., 2006) is widely used for species distribution modeling in ecology, particularly when only presence data is available (West et al., 2016), and has been used in several cases to model risk of plant pathogen invasions (Galdino et al., 2016; Narouei-Khandan et al., 2016). The RF ensemble classification algorithm (Breiman, 2001), is a machine learning method that has also been widely adopted in ecological species distribution modeling due to its high classification accuracy (Bradter et al., 2013; Cutler et al., 2007; Fernández-Delgado et al., 2014; Kampichler et al., 2010). RF is particularly well suited to plant disease incidence datasets which are often non-normal, non-linear, and unbalanced. This method allows for predictors of multiple types (continuous and categorial, for example) and is insensitive to differences in units between predictors (precipitation and solar radiation, for example).

Support vector machines (SVMs) have also previously demonstrated utility for mapping invasive pathogens based on environmental and weather predictors, particularly when data on occurrence and absence is limited (Guo et al., 2005). Although a plethora of machine learning classification algorithms have been developed in recent years, across a wide range of disciplines, RF and SVMs performed best in an evaluation of 179 different classifiers when tested across a range of data sets (Fernández-Delgado et al., 2014). We found that RF was best for classifying our data and it was thus used to estimate environmental risk of establishment in our simulation models.

As is often the case for pathogens emerging in new regions, data for this pathosystem remain sparse and disconnected. This is the crux of the problem in most emerging epidemics: a trade-off between the amount of data to parameterize models for intervention and the ability to act early enough to mitigate spread. This study integrated disparate sources of data to construct a model of pathogen spread and scenario analysis of clean seed provisioning. Disease occurrence and absence data used to train the invasive species distribution model were compiled from different survey sources with different objectives, and often do not represent unbiased systematic observations. Iterative modeling can improve as more data becomes available, while guiding early decision making. Iterative data incorporation can be combined with expert and stakeholder participatory feedback to make the most of available resources.

Decision support for seed system management can potentially be improved by Future considering the potential for economic model improvements. As a cash crop, regional farmer behaviors are often driven by return on investment. For example, stem procurement networks are not static and would rewire based on economics of stem and starch prices. These trends may drive the locations where planting material is needed and sourced, in addition to the number of years a farmer may wait before obtaining off-farm planting material.

Decision support can also be improved with greater epidemiological understanding of *Sri Lankan cassava mosaic virus*. Most current knowledge has been drawn from African cassava mosaic viruses (Holt et al., 1997). High rates of CMD reinfection for clean stems planted under field conditions have been reported in central Cambodia, suggesting that positive and negative selection strategies by farmers may have significant potential in disease management (Malik et al., 2022). Work is ongoing to understand the biotypes of whitefly in the region and their seasonal abundance and dispersal (Götz & Winter, 2016; Ram Kumar et al., 2017; Chi et al, 2020; Leiva et al., 2022). Research is also needed to detect the introduction or recombination of new cassava mosaic virus strains, which may vary in virulence.

Stakeholder feedback during model development can help increase the practical utility of models for specific local decisions. This early feedback can be used to further calibrate models, and allow for data gaps to be identified based on stakeholder opinion. This is particularly important where key data are not systematically published. In this study we implemented a simple stakeholder feedback session. Participatory modeling has been sparsely used in plant disease epidemiology (Gaydos et al., 2019), and more frequently in human epidemiology. There is high value in interactions between modelers and stakeholders, as often the most interesting “modeling questions” do not intersect with the more practical “boots on the ground” issues that need to be addressed. Stakeholder participation provides a meeting ground where these two interests can converge, increasing the potential for real-world use of modeling output.

The development of decision support could also be expanded to include new data about cassava seed systems across the adjacent producing countries of Myanmar and Southern China, which also maintain considerable cassava production. Countries in the insular region of Southeast Asia, including Indonesia and the Philippines, are also at risk for the arrival of CMD, and could benefit from early preventative research on stem exchange patterns and whitefly suitability mapping to identify optimal surveillance and management nodes.

## Acknowledgements

We appreciate the support of the CGIAR Seed Equal Initiative, CGIAR Research Program on Roots, Tubers and Bananas (RTB), and CGIAR Plant Health Initiative, supported by CGIAR Trust Fund contributors (https://www.cgiar.org/funders/). We also appreciate support by USDA NIFA grant 2020-51181-32198. We appreciate helpful comments from E. Goss, R. Muneepeerakul, and I. Small.

## Supplement 1: Full list of environmental predictors

**Table S1.**
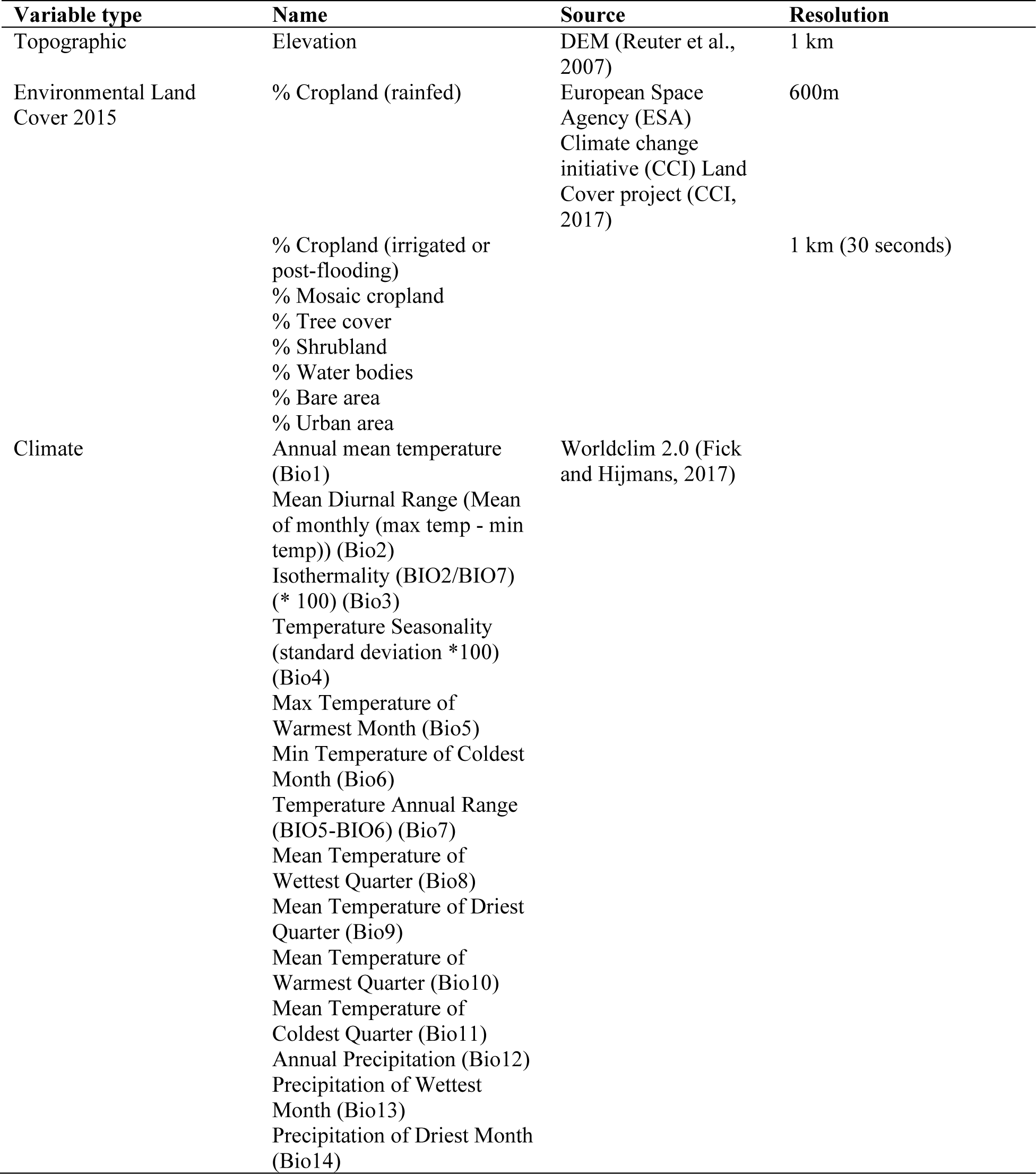
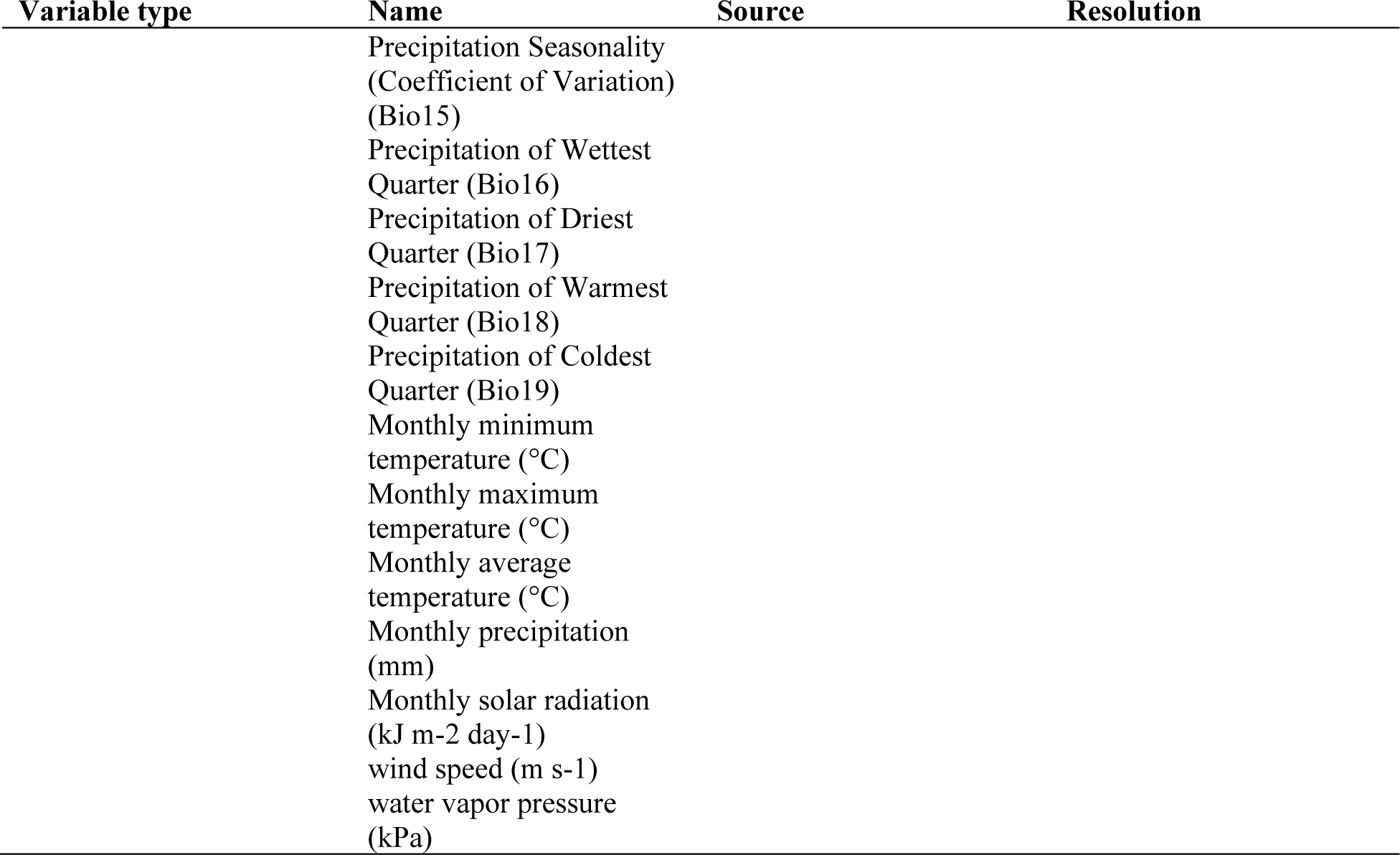
Full list of predictors used in machine learning analysis, prior to dimension reduction step.

## Supplement 2: Workshop survey

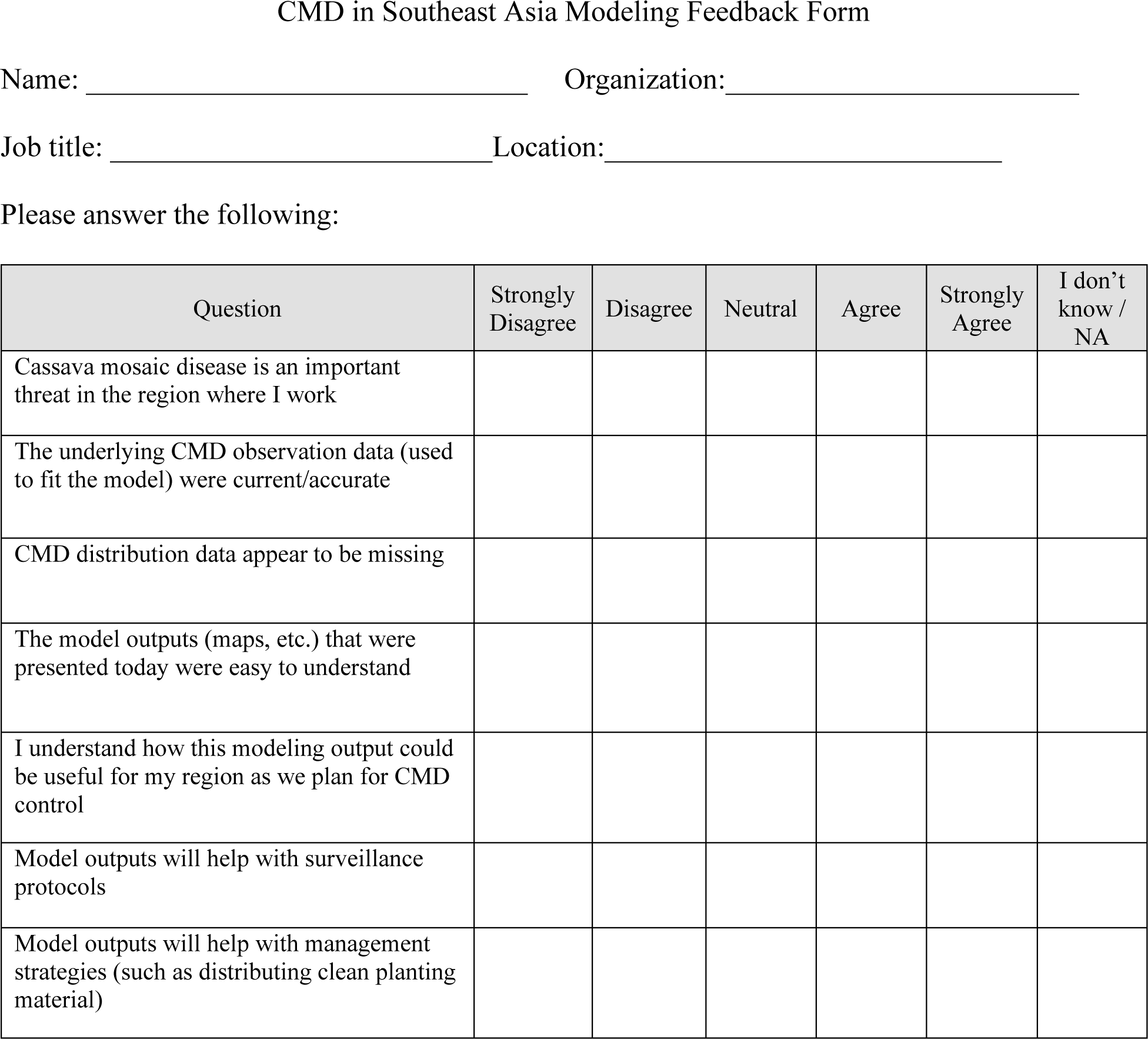

Please describe additional information that you believe should be included in the model. Please describe data CMD observation data that you believe is missing from this analysis. Please describe how you would use the model products / outputs for your work with CMD. How could we change the model output to make it more useful for your needs?

